# Transition of *Vibrio cholerae* through a natural host induces resistance to environmental changes

**DOI:** 10.1101/2021.09.30.462513

**Authors:** Jamie S. Depelteau, Ronald Limpens, Dhrubajyoti Nag, Bjørn E. V. Koch, Jeffrey H. Withey, Annemarie H. Meijer, Ariane Briegel

## Abstract

The pandemic-related strains of *Vibrio cholerae* are known to cause diarrheal disease in animal hosts. These bacteria must overcome rapid changes in their environment, such as the transition from fresh water to the gastrointestinal system of their host. To study the morphological adjustments during environmental transitions, we used zebrafish as a natural host. Using a combination of fluorescent light microscopy, cryogenic electron tomography and serial block face scanning electron microscopy, we studied the structural changes that occur during the infection cycle. We show that the transition from an artificial nutrient-rich environment to a nutrient-poor environment has a dramatic impact on the cell shape, most notably membrane dehiscence. In contrast, excreted bacteria from the host retain a uniform distance between the membranes as well as their vibrioid shape. Inside the intestine, *V. cholerae* cells predominantly colonized the anterior to mid-gut, forming micro-colonies associated with the microvilli as well as within the lumen. The cells retained their vibrioid shape but changed their cell-length depending on their localization. Our results demonstrate dynamic changes in morphological characteristics of *V. cholerae* during the transition between the different environments, and we propose that these structural changes are critical for the pathogen’s ability to colonize host tissues.

## Introduction

*Vibrio cholerae* is a motile, gram-negative bacterium common to fresh and brackish water environments. The pandemic-related strains of *V. cholerae* are known to cause disease in a multitude of animals including humans, fish, and birds (Ali *et al*., 2015; Halpern and Izhaki, 2017; Laviad -Shitrit *et al*., 2017). Upon ingestion by humans, *V. cholerae* colonizes the small intestine utilizing molecular machines such as chemotaxis arrays, flagella, and toxin c-regulated pili to locate and establish its niche (Tacket *et al*., 1998; Butler and Camilli, 2005; Krebs and Taylor, 2011; Utada *et al*., 2014). Within hours of colonization, *V. cholerae* expresses the cholera toxin which causes severe diarrhea in the host. In turn, this releases large quantities of the pathogen back into the environment (Conner *et al*., 2016; Peterson and Gellings, 2018). In humans, *V. cholerae* infection causes over a million Cholera cases and 100,000 deaths annually. Therefore, Cholera remains a public health threat especially in areas of war, natural disasters, and/or poor sanitation (Ali *et al*., 2015).

The journey from its natural habitat, into and through the digestive system, and back into the environment involves extreme changes of the physical and chemical environment. For example, freshwater is considered a sparse environment that contains limited concentrations of salts, minerals, and nutrients (Nelson *et al*., 2008). When ingested, *V. cholerae* must adapt to a sudden change in osmolarity and drop in pH due to the acidity of the stomach (Conner *et al*., 2016). Subsequently, when entering the gut, the bacteria are exposed to bile salts, antimicrobial peptides, and changes in viscosity, all of which are a means of protection against invaders (Merrell *et al*., 2002; Almagro-Moreno *et al*., 2015; Bachmann *et al*., 2015). While these barriers result in the death of some ingested bacteria, *V. cholerae* can adapt and continue its infection cycle by colonizing the intestine, and ultimately re-enter the external environment. How are *V. cholerae* cells able to thrive during these drastic changes in environmental conditions of the infection process? Previous studies have shown that the cells alter their shape in response to changing environments (Bartlett *et al*., 2017; Brenzinger *et al*., 2019; Shi *et al*., 2021). In addition to its shape, *V. cholerae* may also adapt its molecular machines, altering their availability and quantity, to levels that support success in the new environment. For example, the toxin co-regulated pili are generally upregulated as the bacterium approaches the intestinal epithelial cell surface, which aids in attachment and colonization (Krebs and Taylor, 2011; Peterson and Gellings, 2018). However, we currently have limited insight into the detailed morphological changes during the infection cycle of *V. cholerae* inside a host organism.

Here we use cryogenic electron tomography (cryo-ET) and serial block face scanning electron microscopy (SBF SEM) to study the structural adaptations of *V. cholerae* as it transitions through its infection cycle of a natural host, the zebrafish (*Danio rerio*). We chose this model for several reasons: the larvae of the zebrafish are easily accessible for bacterial colonization experiments and microscopy, the physiology of the fish is well understood, and *V. cholerae* has been shown to colonize both adult and larval zebrafish in a manner consistent with human infection and resulting in a diarrheal disease (Runft *et al*., 2014; Murdoch and Rawls, 2019).

Using cryo-ET, we first characterized the 3D architecture of cells grown under nutrient-rich conditions in the laboratory environment and under the nutrient-poor conditions after transition to the freshwater environment of the zebrafish larvae. Next, we compared the architecture of these cells with that of cells that had been excreted back into the environment by the *V. cholerae*-infected zebrafish host. Finally, we used SBF SEM imaging to characterize the localization and cell shape inside the host organism.

Together, our results demonstrate that *V. cholerae* dramatically changes both its shape and the composition of molecular machines in response to this switch of nutrient-rich and nutrient-poor environmental conditions. In contrast, *V. cholerae* within the zebrafish larval intestine, as well as cells excreted from adult and larval zebrafish, maintain their typical comma-like shape during both colonization and excretion back into the environment. These results indicate that the journey through the digestive system of the zebrafish induces physiological and structural changes that protect the bacteria from severe changes in environment, and thus, supports its ability to infect the next host.

## Results

### Environmental *V. cholerae*

For this study, we first characterized *V. cholerae* cells grown in LB media, which is a widely-used nutrient-rich growth condition. *V. cholerae* cells grown overnight in LB exhibit a typical vibroid shape and express the structures associated with bacterial cells in a nutrient-rich environment: chemotaxis arrays, flagella and its motor, uniform spacing between the inner and outer membrane, pili, ribosomes, and storage granules (Fig. 1A). We analyzed the number of cells containing structures related to infection, namely flagella, pili, and the F6 chemotaxis array, as these structures have specific roles in identifying favorable environments and attaching to surfaces. Flagella and F6 chemotaxis arrays were detected in over 80% of bacterial cells and pili in more than 50% (Fig.1B).

**Figure 1:**
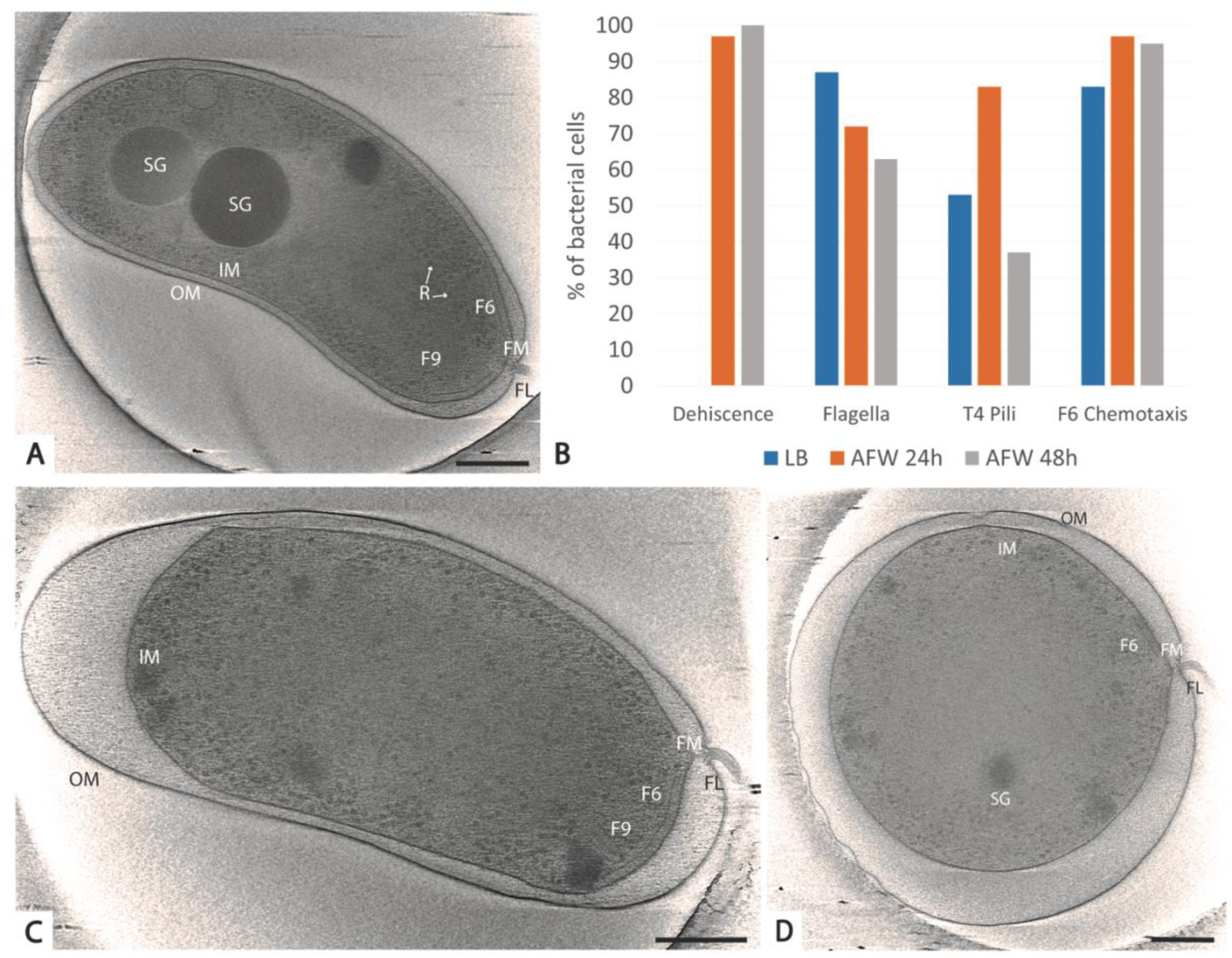
Ultrastructural analysis of V. cholerae transition from LB to artificial fresh water. *V. cholerae* A1552 in LB retain their characteristic comma shape and features (A). Quantification of the presence or absence of the dehiscence phenotype and the molecular machines known to be important in infection are detailed in B. After 24 h in AFW, the shape drastically changes, especially between the inner and outer membrane (C), and this dehiscence phenotype continues after 48 h in AFW (D). OM, outer membrane; IM, inner membrane; SG, storage granule; F6, membrane-imbedded chemotaxis array; FM, flagellar motor; FL, flagella, R, ribosome; F9, cytoplasmic chemotaxis array. Scale bar = 250nm.

We next transitioned the *V. cholerae* grown in LB to artificial fresh water (AFW) and characterized the morphological changes over the course of 48 h. Previous studies have shown that colonization with an El Tor strain peaks at 48 hpi, and thus provided an end point for observation (Runft *et al*., 2014). After 24 h of exposure to AFW, the overall morphology of the cells changed drastically (Figure 1C). While many cells still exhibited an overall vibroid shape, the distance between the inner (IM) and outer (OM) membrane is strongly increased (referred to as dehiscence). Apparently, the inner membrane is constricting while the outer membranes remain the original size. This results in a loose appearance of the OM around the cell and an enlarged periplasmic space. The structures important to the infection cycle, such as the F6 chemotaxis array, the flagella, and the pili were still present in most cells. At 48 h, the morphology changed and dehiscence became the most extreme, while the infection related structures could still be observed in many cells (Figure 1B).

A detailed comparison of the numbers of infection-related structures between the cells grown in LB (n=15), 24 h in AFW (n=29), 48 h in AFW (n=19) shows a decreasing trend in the presence of the flagella over time (87% vs 72% vs 63%), an increasing trend followed by a decrease of the presence of pili (53% vs 83% vs 37%), and a slight increasing trend in the cells with an F6 chemotaxis array (83% vs 97% vs 95%; Figure 1B). Finally, it was also notable to find unknown structures within the cells in AFW for 24 h or 48 h (Figure S1).

### Colonization of the zebrafish larvae intestine

As described in previous research, zebrafish and their larvae are susceptible to *V. cholerae* colonization and the subsequent effects (Runft *et al*., 2014). However, the two fluorescently labeled strains used in this study have a genetic difference that may impact colonization: C6706-tdTomato is deficient in quorum sensing whereas A1552-GFP is quorum proficient. Fluorescence imaging of 5 dpf larvae infected of C6706-tdTomato showed a similar infection pattern as described in Runft et al (which used a quorum proficient strain), and was analogous to the colonization patterns demonstrated in the mouse model using the same strain (Millet *et al*., 2014; Runft *et al*., 2014). In addition, we see significant GFP autofluorescence within the zebrafish larvae, which imposed additional challenges for imaging. Thus, we chose to infect germ-free zebrafish larvae with *V. cholerae* strain C6706-tdTomato to explore the ultrastructural features using serial block face SEM. Similar to the mouse model, colonies were primarily found in the in the intestinal bulb, anterior intestine, and anterior portion of the mid intestine (Figure 2B; Millet et al., 2014) at the junction of several villi, or at or near the base of a single villi. In addition, planktonic cells could be seen swimming within the lumen of the intestinal bulb, implying the environment was conducive to colonization (Figure 2B). This phenotype continued until the mid-gut, which begins approximately just past the swim bladder. At the exit of the intestinal system, the cloaca, large pulses of bacteria could be seen being excreted back into the local environment (Figure 2C).

**Figure 2:**
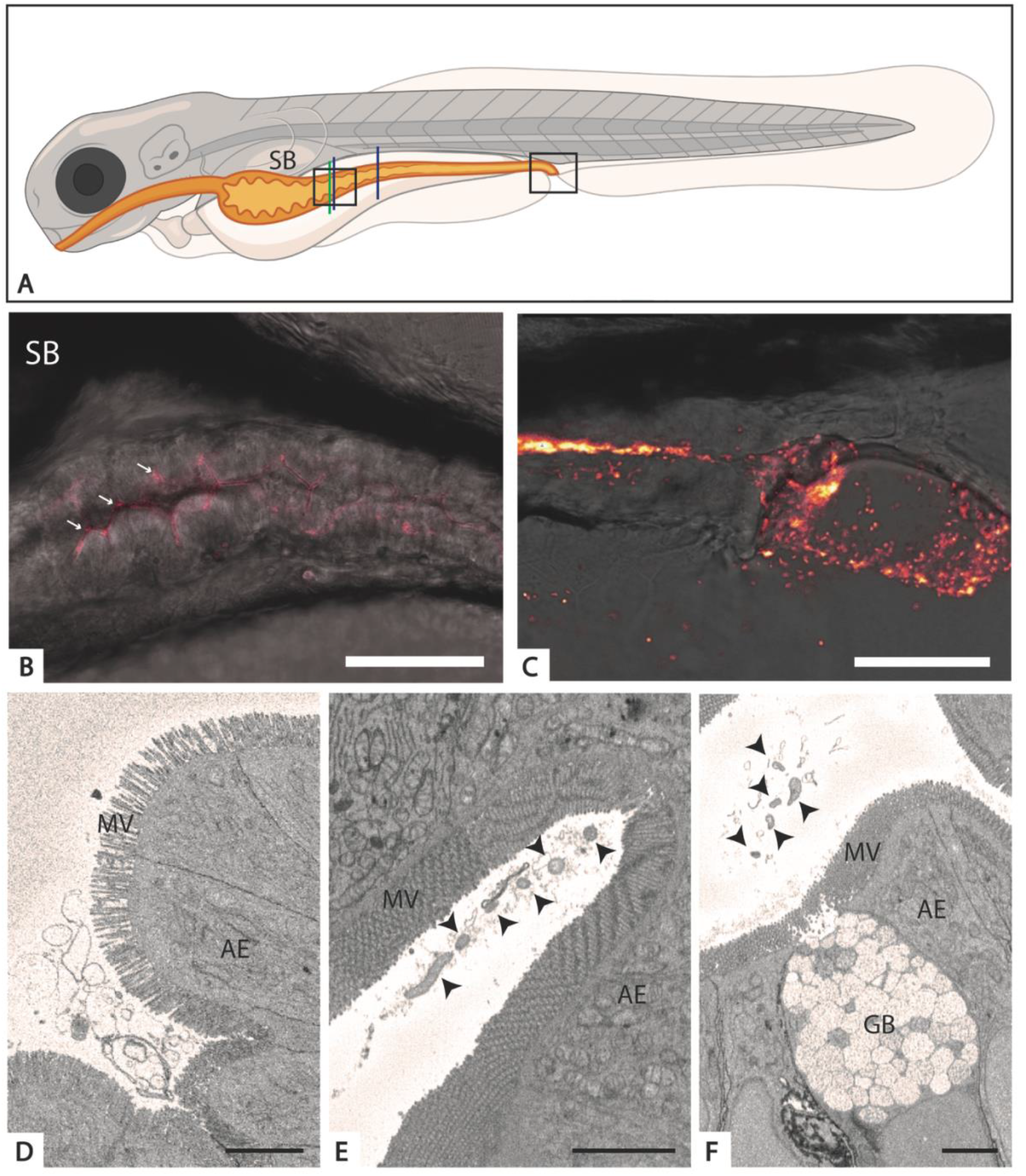
Colonization of the 5 dpf zebrafish larvae with C6706-tdTomato characterized by fluorescence and serial block face scanning electron microscopy. A. A schematic of the zebrafish larvae highlighting the gastrointestinal system (orange) and the areas of interest. B. & C. Fluorescent imaging of the anterior intestine (B, left square in A) and posterior intestine and cloaca (C, right square in A). The white arrows in B indicate areas of colonization. D-F. Representative sections of the germ-free (D, green line in A) and C6706-tdTomato infected zebrafish larvae at the anterior (E, left blue line in A) and mid-intestine (F, right blue line in A).V. cholerae cells are indicated with black arrows. SB, swim bladder; AE, absorptive enterocytes; MV, microvilli; GB, goblet cells. Panel A was created with Biorender.com. Scale bar = 75μm (B); 50μm (C); 3μm (D-F).

Based on this information, we prepared *V. cholerae* C6706-tdTomato infected and uninfected 5 dpf zebrafish larvae for SBF SEM. Using this method, we are able to achieve large volumes of three-dimensional data of areas of interest (in our case covering up to 150 μm length of the intestine). An overview of the germ-free, uninfected zebrafish larvae confirmed the absence of a microbiota, the presence of material in the lumen, and an underdeveloped brush border, as shown by the loose packing of the microvilli (Figure 2D). In contrast, the microvilli of the *V. cholerae*-infected larvae are tightly packed, and microcolonies are present throughout the lumen including the base of villi and unassociated within the lumen (Figure 2E, arrowheads). Similar to the mouse model, colonization of the intestine wanes as it approaches the mid-intestine and posterior intestine, areas where the majority of the mucus producing goblet cells reside (Figure 2F; Millet et al., 2014).

These volumes allowed the analysis of the overall morphology of the bacterial cells that colonized the intestine. Using segmentation software, we were able to outline the bacteria in different parts of the intestine, and subsequently determine the length of the cells (Figure 3A). Overall, all cells exhibited a vibroid shape (Figure 3B). We then determined the length of cells in the villi of the anterior intestine just below the posterior portion of the swim bladder, and the anterior portion of the mid-intestine. A comparison of cell length within each area showed no significant different in cell length (Figure 3C). However, a comparison between the two locations showed a significant difference in cell length (p=0.006; AI-total 1763.05nm vs MI total 1924.10nm): Bacterial cells in the anterior intestine were significantly shorter compared to the cells in the mid-gut.

**Figure 3:**
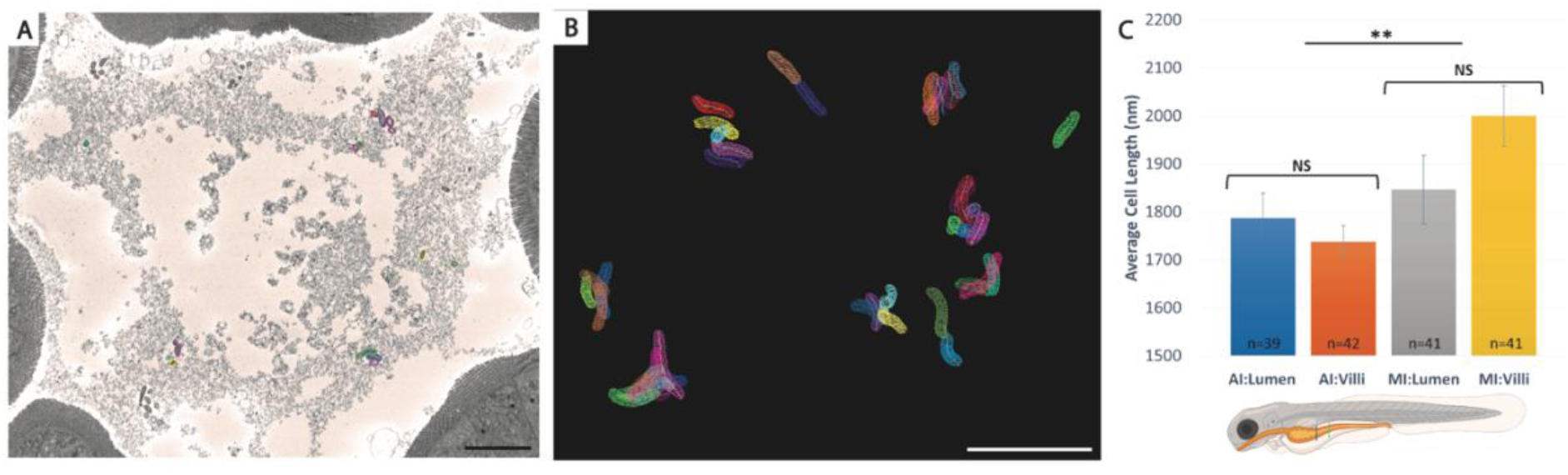
Quantification of V. cholerae cell length at different locations in the zebrafish gut. A. Representative SBF SEM micrograph from the lumen denoting the location of bacteria and highlighting the ones that had been segmented in that section. B. A graphic of most of the bacteria that were segmented for the analysis of bacteria in the lumen and unassociated with the brush border. C. A graph showing the average length of the segmented cells and associated standard deviations. Groups include anterior intestine –lumen (AI-Lumen, blue line), anterior intestine -villi (AI-villi, blue line); mid-intestine lumen (MI-Lumen, green line); mid-intestine-villi (MI-Villi, green line); and the sum of each area: anterior intestine villi and lumen (AI-Total); and mid-intestine villi and lumen (MI-total). Zebrafish schematic in C subset was created in Biorender.com. Scale bars = 5μm. ** p< 0.01.

### Excreted V. cholerae

Finally, we examined the bacteria that are excreted from infected adult and larval zebrafish. Overall, we observed cells in all conditions that had a vibroid or oval shape (Figure 4). None of the groups had a noticeable separation of the inner and outer membrane by 24 h (n=35, 31% from the larvae, n=24; 13% from the adults, compared to nearly 100% of cells in AFW for 24 h or 48 h). For the bacteria that were directly excreted by the adult fish, we observed a similar number of pili (75%) and the F6 chemotaxis arrays (94%) compared to the LB and AFW samples. However, there was a noticeable decrease in the presence of the flagella (10% of cells). The bacteria excreted from the larvae resembled the AFW sample in terms of the presence of the pili (77%) and the F6 chemotaxis arrays (97%), but a noticeable decrease in the presence of the flagella (40% of cells). The presence of storage granules was also noted in the majority of the excreted cells (adult, 88% & larvae, 80% vs. just 31% in the control).

**Figure 4:**
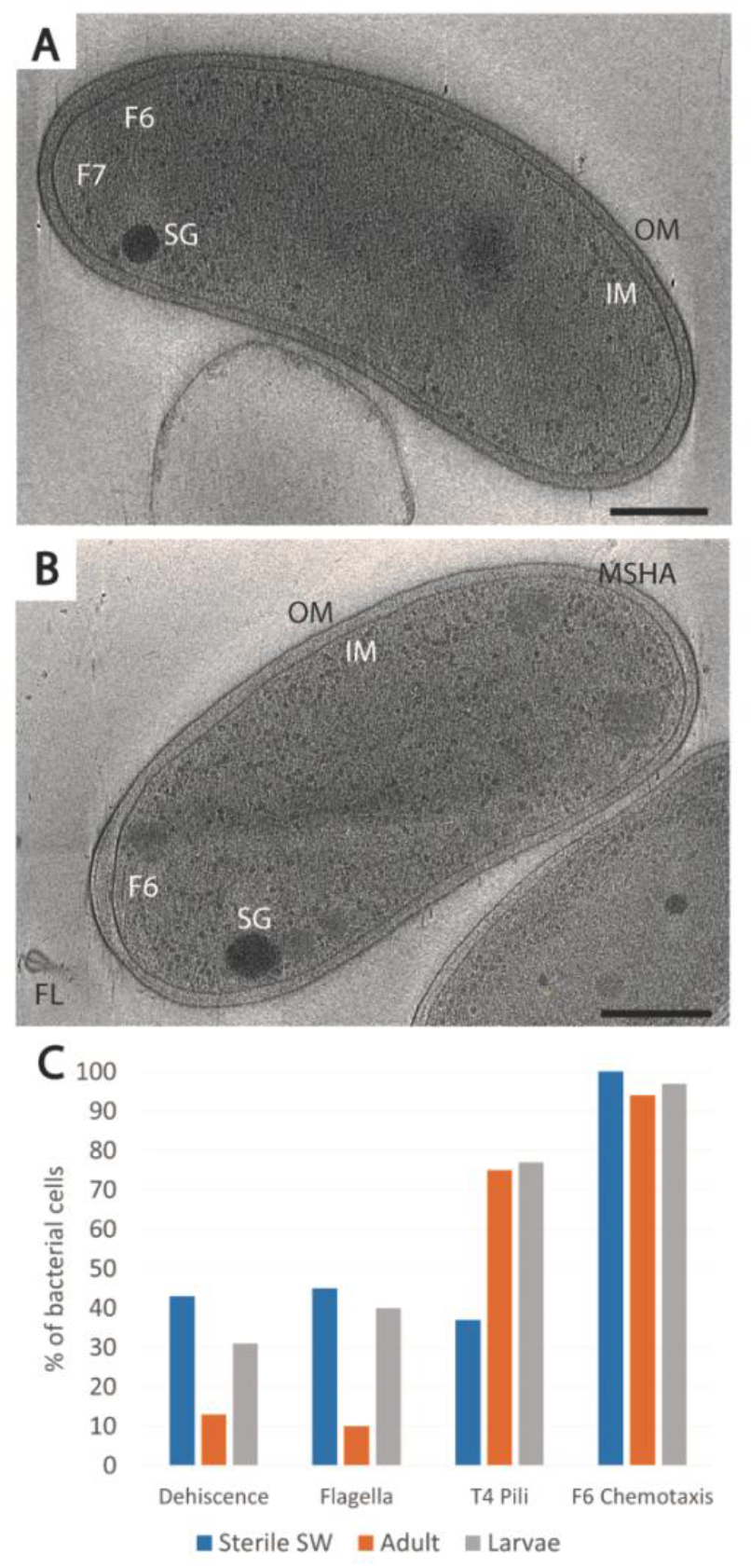
Morphology and machine changes of V. cholerae excreted from zebrafish (adult or larvae). Individual micrographs of a cell excreted from the adult zebrafish (A) or larvae (B). C. Quantification of structures important to the infection cycle. OM, outer membrane; IM, inner membrane; SG, storage granule; F6, membrane-imbedded chemotaxis array; FL, flagella. Scale bar = 250nm.

As a control, we exposed bacteria from the same culture that was used to infect the zebrafish to sterile system water without the presence of fish for 24 h. In this case, we wanted to see what would happen to the bacteria when they were not passaged through the fish but exposed to a similar environment. After 24 h, these cells exhibited dehiscence in about 43% of the population, and the presence of flagella was similar to the adult excreted bacterium (45%; n=49)). A number of cells had a reduction in the number of pili (37%) while the F6 chemotaxis array was present in 100% of the cells (Figure 4). Taken together, these cells do not show as extreme as an affect as seen in cells transition to AFW for 24 h, but seem to be more similar to the cells excreted by the larvae with the exception of the presence of pili.

Finally, as it was noted for the *V. cholerae* isolated from AFW, we also occasionally found unknown structures in the cells excreted from the zebrafish (Figure S1).

## Discussion

The infection cycle of *Vibrio cholerae* involves a complex series of transitions, from fresh or brackish water, into a host’s gastrointestinal system, and then back into the environment. This invariably exposes the pathogen to vastly different environmental conditions, including changes in nutrient availability, pH, and salt concentrations. To what extent the *V. cholerae* cells adjust their ultrastructural morphology to survive and thrive throughout this infection cycle is largely unknown. EM provides an excellent opportunity to explore these changes. The molecular machines of *V. cholerae*, such as the chemotaxis arrays, flagellar motor, and type 6 secretion system, are well characterized and easily identifiable using cryo-ET (Briegel *et al*., 2009; Chen *et al*., 2011; Oikonomou and Jensen, 2016; Rapisarda *et al*., 2019). However, most of these structural analyses were done in *in vitro* conditions and not associated with a natural host. Here, we studied the morphological characteristics of the pathogen during the infection cycle in the natural host model system of the zebrafish.

Using a variety of microscopy techniques, we demonstrate the impact of the environmental changes on the cell morphology of the pathogen at the nanoscale. Using cryo-ET, we show the drastic morphological changes that occur when *V. cholerae* transitions from a high-nutrient and high-salt environment to a low-salt and low-nutrient environment. Upon introduction to artificial fresh water, the bacteria morphologically change, most notably by increasing distance between the inner and the outer membrane. Dehiscence within bacteria, including *V. cholerae*, has previously been described and it has been associated with the transition to viable but non-culturable states, in which cells have a very low metabolic activity (Brenzinger *et al*., 2019; Shi *et al*., 2021). As noted in previous research, *V. cholerae* also expresses unknown structures in response to stressful environments (Dobro *et al*., 2017). In this study, we also observed unknown structures including the previously described ‘wavey filament’ and several types of other filaments (Suppl. Figure S1). It is possible that these structures are related to either nutrient storage or DNA compaction, but further study is needed to clarify their identity and role in the adaption to changes in environmental conditions.

We next examined the bacteria that had transitioned through the zebrafish and been excreted back into a low-salt and low-nutrient environment (sterile system water). Interestingly, unlike the bacteria that were transitioned from LB to AFW, cryo-ET showed that the cells altered their morphology to a lesser extent in response to the environmental change, and a majority retained a vibroid or ovoid shape and displayed MSHA pili and F6 chemotaxis arrays at similar levels to LB-cultured cells. In addition, we observed a decrease in the number of flagella present in the cells excreted from the adult host. This loss of flagella could be the result of flagellar ejection that has been previously observed in response to the transition to a low nutrient environment (Ferreira *et al*., 2019). Together, these results suggest that the transition through the gut of the zebrafish likely primes the bacterium for release into the environment (in our case artificial freshwater or sterilized system water). The majority of the cells retain their cell envelope structure with consistent distance between the IM and the OM. Furthermore, the excreted cells appear to be equipped with the molecular machinery necessary to explore the new environment.

Previous research suggested that *V. cholerae* excreted from the mouse model, as well as cells within or recently dislodged from biofilms, are hyperinfectious, meaning that less cells are needed to continue the infection cycle in another host (Alam *et al*., 2005; Tamayo *et al*., 2010). This has also been found to be true of *V. cholerae* excreted by zebrafish (Nag and Withey, in preparation). Our research supports these findings as the excreted cells did not show typical stress responses such as dehiscence. In addition, they contain molecular machinery, such as the chemotaxis array to sense their environment, the flagella to move towards a favorable environment and MSHA pili to attach to a favorable surface, which are important to effectively transition to another host. Lastly, we also noted that the excreted cells contain storage granules, likely glycogen or polyphosphate, and it was previously shown that these types of storage granules also impart a resistance to the transition from the host back into the environment (von Kruger *et al*., 2006; Bourassa and Camilli, 2009).

Finally, to examine bacteria that had colonized the gut of the zebrafish we focused on the 5 dpf larvae infection model because of its ease of use for imaging and suitability for preparation for serial block face SEM. In our study, we were able to visualize the *V. cholerae* infection of large parts (covering about 150 μm along the length of the larval intestine) at cellular resolution and in three dimensions, which is a substantial improvement in volume compared to previously applied techniques. These data show that the bacteria colonized several parts of the intestine, have the typical vibroid morphology, are actively dividing, and that microcolonies are directly associated with the microvilli or free floating in the lumen. Previous studies suggested that the toxin coregulated pilus (TCP) is involved in this attachment in mammalian models, but the resolution of the SBF SEM is insufficient to confirm or refute this hypothesis at present. In addition, previous research has shown that colonization of the zebrafish model with a *toxT* mutant (which does not produce TCP or cholera toxin) had no colonization defect compared to the wildtype bacterium, indicating that there are other factors involved in colonization (Runft *et al*., 2014). In the future, other methods will be necessary to accurately describe this cell-to-cell interaction and to determine what bacterial appendages play a role in the attachment. Recent developments in large volume sample preparation for cryo-ET are promising options for future studies, but not yet routinely available (Harapin *et al*., 2015; Medeiros *et al*., 2018; Kuba *et al*., 2021).

While our study gives new insights into the morphological characteristics of a *V. cholerae* infection, many questions remain about the lifecycle of this pathogen. In our study, we used a recently described model for infection, and we chose to do this in a germ-free environment to ease the identification of the bacterium in the various microscopy methods. However, understanding our findings in the context of a natural environment will be an essential next step to determine how the intestinal microbiota affects *V. cholerae’s* ability to colonize the host. This is especially true for the understanding how the bacteria interact with the intestinal lining. Current methods do not allow for high resolution imaging at the structural level, as was done with the bacteria in AFW or excreted from the fish.

In summary, we describe the *V. cholerae* infection lifecycle at unprecedented detail using the natural host model, the zebrafish. We show that in laboratory conditions, the transition from nutrient-rich to nutrient-poor conditions causes dramatic morphological changes over 48 h, yet those cells retain factors that are known to be important for infection. We then show that *V. cholerae* excreted from the zebrafish do not show the same changes in morphology, demonstrating that the passage of the bacteria through the intestinal track plays an important role in preparing the bacteria for release back into a freshwater-like environment. Finally, SBF SEM provided insight into the cell morphology and microcolony structure within the lumen. Together, our research provides additional details about the transitions between environments and its role in the infection cycle. Further research will need to be done to confirm physiological mechanism that link structure to function, and this research helps to focus those questions for future studies.

## Experimental Procedures

### Strains

*Vibrio cholerae* El Tor strains A1552, A1552-GFP, and C6706-tdtomato were used throughout this study. A1552 and A1552-GFP are rifampicin resistant (100 μg/ml) strains that were provided by Prof. dr. Melanie Blokesch (École Polytechnique Fédérale de Lausanne, Switzerland). C6706-tdtomato (referred to as VcRed in Millet et al., 2014) is streptomycin resistant (100 μg/ml) and was provided by Prof. dr. Matt Waldor (Harvard University, Boston, MA, USA). Note that this strain carries a mutation in the *luxO* gene rendering the strain quorum deficient. Each strain was stocked in 25% glycerol or 10% DMSO and stored at −80°C until use.

### Animal welfare

All experiments conducted at Leiden University using the ABTL strain were conducted with the approval of the local animal welfare committee (License # 10612) and in accordance with the EU Animal Protective Directive 2010/63/EU. All experiments using the wildtype AB strain were conducted at Wayne State University with the approval of the local IACUC.

Adult fish were maintained according to established protocols published at zfin.org. To generate larvae, on the night before mating, adult fish were separated by sex into small mating tanks containing a removable barrier between the sexes (one female to two males). The following morning, the separation was removed, and the fish were transferred to a medium sized tank containing a mesh-barrier that allows eggs to drop to the bottom of the tank without interference from the adults. Adults were mated in groups of six males to three females per tank for one hour, and then returned to their original housing. Fertilized eggs were collected by pouring the egg-containing system water through a small strainer, washed with system water to remove any debris, and stored in a petri dish until further processing. Eggs generated from the AB strain were incubated at 28°C and washed daily until use. ABTL generated eggs were further processed to render them germ-free.

### Generation of germ-free zebrafish larvae

To generate germ-free larvae, we followed an adapted version of the “natural breeding” protocol previously published (Pham *et al*., 2008). Briefly, freshly laid eggs were collected and immediately transferred in batches of 150-200 eggs to 15 ml tube containing sterile artificial fresh water (sAFW; 60mg/L Instant Ocean; Spectrum Brands, Blacksburg, VA, USA). The sAFW was exchanged for sAFW containing amphotericin B (250 ng/ml), kanamycin (5 μg/ml), and ampicillin (100 μg/ml) and incubated for 4-5h at 28°C. Next, the antibiotic-sAFW was exchanged for sAFW containing 0.1% PVP-I, the eggs were incubated for approximately 45s, and immediately washed three times with sAFW. The sAFW was exchanged with a 0.003% bleach solution and incubated for 5m, then rinsed three times in sAFW. This step was repeated a second time, and in the final wash step the sterile eggs were transferred to a sterile petri dish, unhealthy eggs were removed, and the remaining eggs were stored at 28°C until use. Each day, 100μl of the AFW was plated onto non-selective LB medium to confirm sterility, and then the AFW was exchanged for fresh sAFW.

### Infection model

#### Larvae colonization

Two days before colonization of the larvae, the strain of interest was streaked on selective LB media and incubated overnight at 30°C. The following day, several colonies were collected with a loop and transferred to liquid LB and grown overnight at 30°C with shaking at 180 rpm. On the morning of infection, the OD of the overnight culture was measured and the volume needed for 1e8 bacteria per ml was determined, centrifuged at 5k x g for 5 m, and the resulting pellet was dissolved in 1 ml of sAFW. Simultaneously, the 3-day post fertilization (dpf) zebrafish larvae were separated into batches of 25 larvae in 24 ml of sAFW per petri dish. Subsequently, the 1 ml of bacterial culture was added directly to the liquid. For uninfected larvae, 25 ml of sAFW was added directly to the petri dish. The bacteria were incubated with the larvae for 24h at 28°C, at which point the 4dpf larvae were washed three times in sAFW to remove excess bacteria, and then incubated an additional 24h before processing. For the germ-free ABTL experiments, infection occurred at 3dpf, washing away of excess bacteria at 4 dpf (1 day post infection (dpi)), and imaging or fixation at 5 dpf (2 dpi). For the conventionally reared ZDF larvae experiments, the same protocol was used, but infection occurred at 5 dpf and processing was done at 7 dpf (2 dpi).

#### Adult

Adult, conventionally reared AB wildtype zebrafish were transferred to a 500 ml beaker filled with system water and fasted for 12 h before infection. On the day of infection, the water was exchanged for sterile system water and subsequently inoculated with *V. cholerae*. As with the larvae, bacteria were first streaked on a selective LB plate, followed by the growing of an overnight culture. An adequate volume of overnight culture to generate a final concentration of 2.7e7 bacteria/ml was centrifuged as above, the pellet resuspended in sterile system water, and then added directly to the beaker. After 24 h of infection, the fish were washed in sterile system water two times, fresh sterile system water was added, and then left for another 24 h. The adult fish were sacrificed for other experiments. The supernatant was collected, centrifuged at 8k x g for 10m, and most of the supernatant was carefully removed. The remaining supernatant was used to wash the sides of the tube and redissolve the pellet, and then transferred to an 1.5 ml tube. 1.5 μl of anti-*V. cholerae* antibody was added, incubated for 5min, and then centrifuged again for 10m at 8k x g. The supernatant was carefully removed, the pellet was dissolved sterile system water and 5nm gold beads were added, and the sample immediately used for plunge freezing.

### Fluorescence imaging

Fluorescence imaging occurred using a Leica TCS SPE inverted confocal scanning microscope (Leica Microsystems, Wetzlar, Germany). 5 dpf zebrafish larvae were first anesthetized in 0.2 mg/ml tricaine, and subsequently mounted laterally in a drop of 1.3% low melting agarose on a Willco-dish glass bottom microscopy dish (Willco Wells B.V., Amsterdam, The Netherlands). Once the agarose solidified, the fish were surrounded by 0.2 mg/ml tricaine and then imaged using the 40x long working distance water immersion lens. Z-stacks were acquired using the Leiden Application Suite X (LAS X).

### Sample collection and preparation

Samples involving bacteria incubated in LB and AFW were prepared as follows. Bacteria were prepared as described for the infection model. 1e^8^ bacteria per ml were isolated from an overnight culture, pelleted, and redissolved in 1 ml of sAFW. Subsequently, this was added to a sterile petri containing 24 ml of sAFW with or without zebrafish larvae and incubated at 28°C. 2 ml of the diluted bacteria was collected at 1dpi and 2dpi, centrifuged for 5m at 5k x g, and the resulting pellet was dissolved in 27 μl sAFW and 3 μl 10 nm gold beads (Cell Microscopy Core, Utrecht University, Utrecht, The Netherlands). For each time point, grids were prepared using a Leica EM GP (Leica Microsystems, Wetzlar, Germany) by adding 3 μl of the sample directly to a glow discharged Quantifoil R2/2, 200 mesh Cu grid (Quantifoil Micro Tools GmbH, Jena, Germany), incubated for 30s, blotted for 1s and immediately plunged into a liquid ethane cooled to −184°C. Grids were transferred to grid boxes and stored in liquid nitrogen until imaging.

Samples associated with the AB zebrafish and larvae were prepared using a portable manual plunger (Depelteau *et al*., 2020). In brief, 3 μl of sample was added to a glow discharged Quantifoil R2/2, 200 mesh grid, incubated for 30s, and manually plunged into a liquid ethane/propane mixture cooled by liquid nitrogen (Tivol *et al*., 2008). For experiments using germ-full larvae and adults, the anti-*Vibrio cholerae* polyclonal antibody (KPL Bactrace, ELITechGroup, Spankeren, Netherlands) was added to the excreted bacteria sample as a way to identify *V. cholerae* cells during target selection for cryo-ET. The addition of the 5 nm beads during the sample preparation process acted as a secondary that was easily visualized in the electron microscope.

### Cryo-EM

All cryo-EM samples were screened using a Thermo Fisher Scientific (TFS; Waltham, MA, USA) Talos L120C equipped with a Ceta CMOS camera and extended cooling. Samples were inserted using a Gatan 626 side entry holder (Gatan Inc, Pleasanton, CA, USA).

Cryogenic electron tomography data was collected using a TFS Titan Krios microscope equipped with either a Gatan K2 BioQuantum or the Gatan K3 BioQuantum direct electron detection camera, both equipped with a post-column energy filter operating with a slit width of 20eV. The data collected from cells in AFW was collected using UCSF Tomography using a bidirectional tilt scheme of −54° to 54° with 2° increments, the K2 camera with a pixel size of 5.44Å, a defocus of −6μm and total dose of 100 e^-^/Å^2^ (Zheng *et al*., 2007) The data collected from excreted cells was collected using SerialEM with a bidirectional tilt scheme of −54° to 54° with 2° increments, the K3 camera with a pixel size of 4.41Å. a defocus of −8um, and total dose of 140 e^-^/Å^2^ (Mastronarde, 2005).

### Serial Block Face Scanning Electron Microscopy

After fixing the material for 2 h at room temperature with 2.5% GA + 2% PFA in 0.15 M Cacodylate buffer containing 2 mM CaCl_2_ , the material was washed 3 times with buffer and then placed into 2% OsO_4_ / 1.5% potassium ferrocyanide in 0.15 M Cacodylate buffer containing 2 mM CaCl_2_. The material was left for 60 minutes on ice. After washing 3 times in milliQ water, the material was placed into 1% thiocarbohydrazide for 20 m at room temperature. The material was again washed and then stained with 2% aqueous OsO_4_ for 30 m at room temperature. After washing 3 times, the material was placed into 1% Uranyl acetate for 2 h at room temperature. The material was washed with milliQ water then stained with lead aspartate for 30 m at 60°C. The material was again washed with milliQ water and then dehydrated on ice in 20%, 50% and 70% ethanol solutions for 5 m at each step. After replacing the 70% ethanol with a fresh 70% ethanol solution, the samples were kept overnight at 4°C. The next day, samples were dehydrated in 90%, 100%, 100% ethanol solutions for 5 m at each step. Next, the material was kept in dry acetone for 10 m on ice, and another 10 m in fresh dry acetone at room temperature. The material was infiltrated with 25%, 50% and 75% Durcupan ACM solution in acetone for 2 h at room temperature for each step, followed by an overnight step at room temperature in 100% Durcupan resin. The next day, the material was placed in fresh Durcupan resin for 2 h at room temperature, after which the material was flat embedded and polymerized at 60°C for 48 h.

Data were collected with a 3View2XP (Gatan Inc, Pleasanton, CA, USA) unit installed on a Zeiss Gemini 300 field emission SEM (Carl Zeiss Microscopy GmbH, Jena, Germany). The volumes were collected at 1.8 kV accelerating voltage and variable pressure at 5 Pascal. The pixel dwell time was 2 μs, with a pixel size of 10 nm and a section thickness of 75 nm.

### Imaging processing and statistical analysis

Imaging data collected from the cryo-electron microscope was processed using the IMOD image processing suite (Kremer *et al*., 1996). Initially, frames were aligned using the alignframes feature, and the resulting tilt series were batch processed using batchruntomo (Mastronarde and Held, 2017). The initial tomograms were reviewed for quality, and select tomograms were then further processed to improve the bead model and positioning, and finally reprocessed with a SIRT-like filtered back projection to improve the contrast.

Segmentation and cell length measurements were obtained using the 3dmod component of the IMOD image processing suite (Kremer *et al*., 1996). Length measurements were normally distributed, variance was compared and then significance was determined using a pool-variance two sample T-test (Statistics Kingdom, 2021).

## Acknowledgements

The authors would like to thank Prof.dr. Melanie Ohi and Louise Chang of the University of Michigan for the use of their glow discharger during sample collection at Wayne State University, and the Netherlands Center for Electron Nanoscopy and its operators for assistance during data collection.

## Supplementary Figure

**Figure S1.**
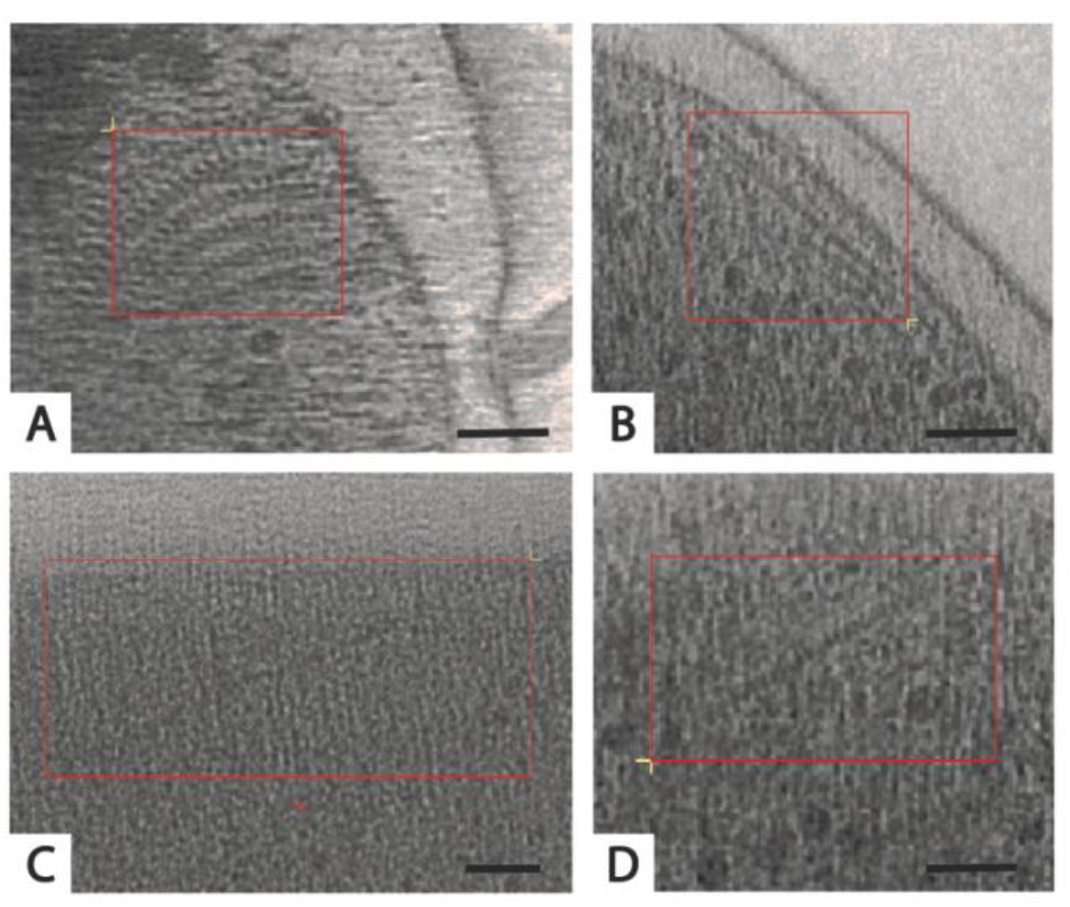
Unknown structures observed in cells in freshwater environments. A. Wavey filament. B. Linear filament 1. C. Periplasmic filament. D. Linear filament 2. Red box identifies boundaries of structure. Scale = 50nm

